# Elevated prevalence of azole resistant *Aspergillus fumigatus* in urban versus rural environments in the United Kingdom

**DOI:** 10.1101/598961

**Authors:** Thomas R Sewell, Yuyi Zhang, Amelie P Brackin, Jennifer MG Shelton, Johanna Rhodes, Matthew C Fisher

**Affiliations:** MRC Centre for Global Infectious Disease Analysis, Department of Infectious Disease Epidemiology, Imperial College London, London, UK

**Keywords:** Aspergillus fumigatus, triazoles, antifungal resistance, fungal pathogen, fungal epidemiology, multidrug resistance, environment

## Abstract

Azole resistance in the opportunistic pathogen *Aspergillus fumigatus* is increasing, dominated primarily by two environmentally-associated resistance alleles: TR_34_/L98H and TR_46_/Y121F/T289A. Using an environmental sampling strategy across the South of England we assess the prevalence of azole resistant *A. fumigatus* (AR*Af*) in soil samples collected in both urban and rural locations. We characterise the susceptibility profiles of the resistant isolates to three medical azoles, identify the underlying genetic basis of resistance and investigate their genetic relationships. AR*Af* was detected in 6.7% of the soil samples, with a higher prevalence in urban (13.8%) compared to rural (1.1%) locations. Nineteen isolates were confirmed to exhibit clinical breakpoints for resistance to at least one of three medical azoles, with 18 isolates exhibiting resistance to itraconazole, four to voriconazole, with two also showing additional elevated minimum inhibitory concentration to posaconazole. Thirteen of the resistant isolates harboured the TR_34_/L98H resistance allele and six isolates carried TR_46_/Y121F/T289A allele. The 19 azole-resistant isolates were spread across five *csp1* genetic subtypes, t01, t02, t04B, t09 and t18 with t02 the predominant subtype. Our study demonstrates that AR*Af* can be easily isolated in the South of England, especially in urban city centres, which appear to play an important role in the epidemiology of environmentally-linked drug resistant *A. fumigatus*.

## Introduction

*Aspergillus fumigatus* is a ubiquitous ascomycete fungus with a pan-global population distribution and a primary ecological niche of decaying vegetation and soil (1). This fungus is the most prevalent species among ~250 described Aspergilli, partly due to its ability to survive and grow in a wide range of conditions, but also through the large-scale dispersal of airborne conidia (1–3). *A. fumigatus* is also an opportunistic pathogen and is commonly responsible for aspergillosis, a spectrum of clinical syndromes caused by *Aspergillus* spp. that affects millions of individuals worldwide (4). Invasive aspergillosis (IA), the most severe form of the disease, can lead to serious and even fatal illness in immunocompromised individuals, with a mortality rate of 40 - 90% (5, 6).

Triazole antifungals are used for the treatment and prophylaxis of *Aspergillus* spp. infections, although resistance has emerged, often conferred by the presence of mutations in lanosterol 14 alpha demethylase (erg11, syn. CYP51), which is a key component of the ergosterol biosynthetic pathway and target for azole antifungals (7, 8). Recently, azole-resistant *A. fumigatus* (AR*Af*) has emerged environmentally, where selection is thought to be driven by the broad application of agricultural azole fungicides; structurally similar to their medical counterparts and indistinguishable in their mode of action (9, 10). Environmentally sourced AR*Af* are typically found with a tandem repeat mutation in the *cyp51A* promoter region and linked single nucleotide polymorphisms in the coding region, with common examples being TR_34_/L98H and TR_46_/Y121F/T289A (11, 12).

Studies have shown that AR*Af* isolates harbouring resistance-associated *cyp51A* variants are globally distributed, and are often found alongside wild-type *A. fumigatus* in a diverse set of environmental substrates, including: agricultural soil (13, 14), flower beds (13, 15–17) and timber mills (18). Moreover, clinical cases of aspergillosis with AR*Af* isolates harbouring either TR_34_/L98H or TR_46_/Y121F/T289A continue to emerge (19, 20), with one study specifically linking a fatal case of aspergillosis to a genotypically indistinguishable isolate sourced from the patients own home (21). Retrospective studies of patients with invasive aspergillosis (IA) and infected with azole-resistant genotypes of *A. fumigatus* show an excess mortality of 25% at day 90 when compared against patients with wild-type-infections (22).

Despite a generally good understanding of AR*Af* global prevalence, very few studies have investigated resistance in the United Kingdom or focused sampling strategies across a diverse set of substrates. In this study, we aim to determine the prevalence of AR*Af* in soil collected from a wide range of sites across the UK’s southernmost region. Sampling locations include ancient woodlands, agricultural fields, tourist attractions and densely populated city centres. We show that the UK has similar prevalence rates to other countries and that both TR_34_/L98H and TR_46_/Y121F/T289A can be regularly isolated from urbanised locations, especially flower beds in close proximity to city centre hospitals.

## Materials and Methods

### Environmental sampling

Samples were collected from 16 sites across South England between May and July 2018. Locations were selected to include a range of habitat types and included remote forested regions, urban city centre, agricultural and flower fields. At each sample site, dry surface soil was loosened and collected into a 5ml Eppendorf tube (Eppendorf AG, Hamburg, Germany). All tubes were labelled with site description, longitude and latitude coordinates. All samples were stored at 4°C until processing.

### Recovering of *A. fumigatus* and screening for azole resistance

To isolate *A. fumigatus* from environmental samples 2 g of each sample was suspended in 8 ml of sterile distilled water with 0.85% NaCl (SIGMA, Sigma-Aldrich Chemie Gmbh, Steinheim, Germany) plus 0.01% Tween 20 (SIGMA, Sigma-Aldrich Chemie Gmbh, Steinheim, Germany) and vortexed vigorously for 1 min. After 1 min settling, 200 μl of supernatant was added to two control plates, containing sabouraud dextrose agar (CM0041, Oxoid Ltd, Basingstoke, Hants, UK) supplemented with 16 g/ml penicillin G sodium salt (SIGMA, Sigma-Aldrich Chemie Gmbh, Steinheim, Germany) and 16 g/ml streptomycin sulfate salt (SIGMA, Sigma-Aldrich Chemie Gmbh, Steinheim, Germany), and two azole-containing plates containing sabouraud dextrose agar supplemented with 16 g/ml penicillin G sodium salt, 16 g/ml streptomycin sulfate salt and 8 g/ml tebuconazole (PESTANAL analysis standard, Sigma-Aldrich AG Industriestrasse, Switzerland). These plates were incubated at 42°C and examined for growth after 72 hours. *A. fumigatus* isolates were identified by observation of their macro and microscopic morphology.

### *In vitro* susceptibility testing - Minimal inhibitory concentration (MIC) measurement

Seven wild-type isolates and 19 suspected azole-resistant isolates were tested for antifungal drug susceptibility against three medical azoles, itraconazole (ITC), voriconazole (VRC) and posaconazole (PSC) using the European Committee Antimicrobial Susceptibility Testing (EUCAST) microdilution method using MICRONAUT-AM EUCAST MIC (Minimum Inhibitory Concentration) plates (Merlin Diagnostika GmbH, Bornheim, Germany). The reference wild-type clinical *A. fumigatus* isolates Af293 and an environmental azole-resistant isolate BUU09 (TR_34_/L98H) were used as control strains. All isolates were grown for 72 hours at 37 °C before washing with 8 to 10 ml of 0.01% Tween 20. The resulting conidial suspension was filtered through Whatman 1 filter paper (Whatman plc, GE Healthcare Life Sciences, UK) to remove hyphae and debris. The filtered conidial suspension was adjusted to 0.5 McFarland with phosphate-buffered saline (PBS pH7.4 (1X), gibco, Life Technologies Limited, Paisely, UK) using a BioTek ELx808 spectrophotometer (BioTek Instruments Ltd, Winooski, USA). The adjusted conidial suspension (0.5 ml) was added to 9.5 ml of the MICRONAUT-RPMI medium (MICRONAUT-RPMI Medium + MOPS + Glucose, Merlin Diagnostika GmbH, Bornheim, Germany) and 100 μl of the mixture was then used to populate the MICRONAUT-AM test plate. All plates were examined after 48-hour incubation at 37°C in a humid chamber. The MIC endpoint was read visually and determined as the lowest concentrations of the azoles yielding no visible growth. The results of all susceptibility tests were interpreted using EUCAST clinical breakpoints [91]. The isolates were regarded as susceptible when the MIC was ≤ 1 mg/l for ITC and VRC, and ≤ 0.125 mg/l for POS; and resistant when the MIC was > 2mg/l for ITC and VRC, and > 0.25 mg/l for POS. Isolates with MIC values in between the susceptible and resistant breakpoint were considered to be intermediately susceptible.

### DNA extraction for sequence analysis

Genomic DNA was extracted using a modified MasterPure Yeast DNA Purification (Lucigen Corporation, Cambridge, UK) protocol which included an additional bead-beating treatment to enhance DNA yield. Cultured *A. fumigatus* isolates were washed in 10 ml of 0.05% Tween 20; 1.8 ml of the conidial suspension was transferred into a 2 ml Eppendorf tube and centrifuged at 8000 rpm for 5 mins. The condial pellet was mixed with 300 μl of Yeast Cell Lysis Solution and 1 μl of 5 g/l RNase A and transferred into a new 2 ml Eppendorf tube containing 1.0 mm diameter Zirconia/Silica Bead (Thistle Scientific, Glasgow, UK). The tube was subjected to bead beating for 3 x 45 seconds at 30 m/s in a TissueLyser II (Qiagen, Hilden, Germany) and placed on ice for 2 mins before repeating. The lysate was transferred to a new 1.5 Eppendorf tube and was twice centrifuged at 13000 rpm for 2 mins to pellet debris. The supernatant was transferred to a 1.5 Eppendorf tube and incubated at 65°C for 15 mins before placing on ice for another 30 mins. After incubation, 150 μl of MPC Protein Precipitation Reagent was added to the sample and mixed by pulse vortexing for 10 seconds. For further pelleting of cellular debris, the mixture was centrifuged at 13000 rpm for 2 mins and the supernatant was transferred to a clean 1.5 Eppendorf tube. DNA was precipitated by adding 500 μl of isopropanol (SIGMA, Sigma-Aldrich Chemie Gmbh, Steinheim, Germany) to the supernatant before 4°C centrifugation at 14000 rpm for 10 mins. The supernatant was removed, and the remaining DNA pellet washed with 500 μl of 70% ethanol (SIGMA, Sigma-Aldrich Chemie Gmbh, Steinheim, Germany) before centrifugation at 13000 rpm for 2 mins. After removing the ethanol, the DNA pellet was air-dried for approximately 10-15 mins in a biosafety cabinet. Finally, 40 μl of nuclease-free water was added to the DNA pellet and left to resuspend at 4°C for 24 hours. The DNA was either used immediately or stored at −20°C until required. DNA yield was quantified using a Qubit 2.0 Fluorometer (Invitrogen by Thermo Fisher Scientific corporation, Massachusetts, USA) and the Qubit dsDNA BR (Broad-Range) Assay Kit (Invitrogen) according to the products guidelines.

### PCR amplification

PCR was used to amplify part of the *cyp51A* gene and promoter, the *csp1* region and *beta-tubulin* as previously described [45], [80]. Amplification of *cyp51A* was performed using the L98HR primer (5’-TTCGGTGAATCGCGCAGATAGTCC-3’) and TR34R primer (5’-AGCAAGGGAGAAGGAAAGAAGCACT-3’) (Invitrogen) at 100 nM, 20.125 μl nuclease-free water (Qiagen, Hilden, Germany), 2.5 μl PCR Buffer (Qiagen PCR Buffer containing 15mM MgCl_2_, Qiagen, Hilden, Germany), 0.5 μl dNTP mix (dNTP Set, PCR Grade, 10mM each, Qiagen, Hilden, Germany), 0.125 μl Taq DNA Polymerase (5 units/L, Qiagen, Hilden, Germany) and 1 μl of DNA template. Amplification conditions were: a 2 mins denaturation step at 94°C, followed by 35 cycles of 15 seconds at 94°C, 30 seconds at 57°C, and 30 seconds at 72°C. A final elongation step for 2 mins at 72°C was followed after the last cycle. Amplification of *csp1* was performed using the CSP1F primer (5’-TTGGGTGGCATTGTGCCAA-3’) and CSP1R primer (5’- GAGCATGACAACCCAGATACCA-3’) at 100 nM, 20.125 μl nuclease-free water, 2.5 μl PCR Buffer, 0.5 μl dNTP mix, 0.125 μl Taq DNA Polymerase and 1 μl of DNA template. Amplification conditions were: a 5 mins denaturation step at 94°C, followed by 35 cycles of 15 seconds at 94°C, 30 seconds at 55°C, and 30 seconds at 68°C. A final elongation step for 2 mins at 68°C was followed after the last cycle. Amplification of *beta-tubulin* was performed using the Bt2A_F primer (5’-GGTAACCAAATCGGTGCTGCTTTC - 3’) and Bt2A-R primer (5’ - ACCCTCAGTGTAGTGACCCTTGGC - 3’) at 100 nM, 20.125 μl nuclease-free water, 2.5 μl PCR Buffer, 0.5 μl dNTP mix, 0.125 μl Taq DNA Polymerase and 1 μl of DNA template. Amplification conditions were: a 5 mins denaturation step at 95°C, followed by 35 cycles of 30 seconds at 95°C, 30 seconds at 55°C, and 1 min at 72°C. A final elongation step for 5 mins at 72°C was followed after the last cycle.

### PCR product visualisation, purification and sequencing

Both *cyp51A* gene and *csp1* gene PCR products were visualised on a 2% agarose gel using agarose gel electrophoresis. 100 ml of 2% agarose solution was prepared by mixing 2 g of agarose powder with 100 ml of TBE buffer. 10 μl of SafeView Nucleic Acid Stain was added for visualisation of the PCR product in the gel. Before loading the samples and molecular weight ladder (Quick-load Purple 50 bp DNA Ladder and Quick-load Purple 1 kb DNA Ladder, New England Biolabs Inc, UK) to the gel, 1 μl of Gel Loading Dye was mixed with 5 μl of each sample. Finally, the gel was run at 120 to 130 V for approximately 45 to 60 mins. The DNA fragments were visualised using a G:Box Gel Image Analysis System (Syngene UK, Cambridge, UK). The PCR products were then purified using ExoSAP-IT PCR Product Cleanup Reagent (ThermoFisher Scientific, Massachusetts, USA) following the product guidelines. Briefly, 7.5 μl of each post-PCR reaction product was mixed with 3 μl of the reagent. The mixture was incubated for 15 mins at 37°C in order to degrade the remaining nucleotides and primers in the PCR product, followed by incubation of 15 mins at 80°C to inactivate the reagent. The treated products were then stored at −20°C until required. The treated PCR products and sequencing primers, L98HR, TR34R, CSP1F and CSP1R were prepared for sequencing following the DNA Sanger sequencing sample submission guidelines provided by GENEWIZ (GENEWIZ UK LTD, Takeley, UK). They were then sent away for Sanger sequencing by GENEWIZ UK laboratory.

### Sanger sequencing of *cyp51A* mutations, *csp1* types and *beta-tubulin*

Sanger sequencing results from all selected isolates were trimmed and assembled using CLC Main Workbench 8.0.1 software (QIAGEN Bioinformatics, Hilden, Germany), with alignment stringency set to medium. The forward and reverse reads were merged into a consensus sequence using the assemble sequences function. The consensus *cyp51A* sequences of each environmental isolates, and the known TR_34_/L98H isolate BUU09, were aligned against the referenced wild-type isolate, *A. fumigatus* Af293. The isolates that exhibited the same 34-bp tandem repeat as the isolate BUU09 were confirmed to harbor the TR_34_/L98H allele. The isolates that had a 46-bp tandem repeat were compared with *A. fumigatus* isolate VRJ056 (GenBank accession number: MF070884.1), a known clinical isolate harboring the TR_46_/Y121F/T289A mutation, to confirm their resistant mechanism. The *csp1* types of each environmental isolates and two control isolates were assigned according to the *csp1* typing nomenclature described previously (23), which was based on manually inspecting the repeat regions in each consensus *csp1* sequences.

## Results

### Environmental *A. fumigatus* isolates

A total of 178 soil samples were collected across Southern England, including soil and compost samples obtained from public gardens, parks, cemeteries, and flower beds outside hospitals. Samples were collected in Central London (n = 64), Bath (n = 8), flower beds around Stonehenge (n = 8), remote forested regions in the New Forest National Park (n = 46), a lavender farm in Surry (n = 13) and farmland in Cambridgeshire (n = 39). Of the 178 soil samples, 131 (74%) were positive for *A. fumigatus* type growth on control plates, with a varied recovery rate amongst different sampling sites (Table 1). The highest recovery rate was from Central London (93.8%) and the lowest from Cambridgeshire (35.9%). Eleven *A. fumigatus* positive soil samples yielded a total of 83 *A. fumigatus* type isolates that were able to grow on the azole-containing plates. Nine of these soil samples were collected from urban locations in London (n=8) and Bath (n=1). Among the eight samples from London, four were collected from flower beds outside Royal Free Hospital and The Whittington Hospital, and another two samples were obtained from compost in Waterlow Park, 500 meters away from The Whittington Hospital. Nineteen putatively resistant isolates were selected for further MIC testing and sequence analysis, ensuring representation from each azole positive soil samples. Seven control isolates were randomly selected for further analyses, representing each sampling site. All 26 isolates were identified as *A. fumigatus* by sequencing of the *beta-tubulin* gene.

**Table 1.** Soil sampling areas and *A. fumigatus* recovery rate, with azole resistance determined by MIC and *cyp51A* sequencing.

| Sampling site | Samples collected (n) | Number of samples with *A. fumigatus* growth (%) | Number of samples recovered azole-resistant *A. fumigatus*<sup>a</sup> | Prevalence of azole-resistant *A. fumigatus* |
| --- | --- | --- | --- | --- |
| **London** |  |  |  |  |
| Overall | 64 | 60 (93.8%) | 9 | 14.0% |
| Kensal Green Cemetery | 5 | 5 (100%) | 0 | 0% |
| Queen Mary's Rose Garden & Regents Park | 8 | 7 (87.5%) | 0 | 0% |
| The Whittington Hospital & Waterlow Park | 5 | 5 (100%) | 3 | 60.0% |
| Hampstead Heath | 3 | 1 (33.3%) | 0 | 0% |
| Royal Free Hospital | 5 | 5 (100%) | 3 | 60.0% |
| Green Park & Hyde Park | 9 | 9 (100%) | 1 | 11.1% |
| Charing Cross Hospital & Margravine Cemetery | 8 | 7 (87.5%) | 0 | 0% |
| Brompton Cemetery | 6 | 6 (100%) | 1 | 16.7% |
| Elstree Open Space | 8 | 8 (100%) | 1 | 12.5% |
| Flower beds in urban city | 7 | 7 (100%) | 0 | 0% |
| **Bath city centre, Somerset** |  |  |  |  |
| Flower beds | 8 | 4 (50%) | 1 | 12.5% |
| **Stonehenge, Wiltshire** |  |  |  |  |
| Flower beds and farm | 8 | 6 (75%) | 1 | 12.5% |
| **New Forest National Park, Hampshire** |  |  |  |  |
| Remote forest | 46 | 37 (80.4%) | 1 | 2.2% |
| **Surrey** |  |  |  |  |
| Lavender farm | 13 | 10 (76.9%) | 0 | 0% |
| **Cambridgeshire** |  |  |  |  |
| Wheat farm | 30 | 12 (40%) | 0 | 0% |
| Open space surrounded by farm | 9 | 2 (22.2%) | 0 | 0% |
| COMBINED TOTAL | 178 | 131 (73.6%) | 12 | 6.7% |

a. Resistance determined by both MIC testing and sequencing analysis.

### MIC measurement

All 19 azole-tolerant isolates were confirmed to be resistant to at least one of the three medical azoles tested (Table 2). Eleven of these exhibited resistance to itraconazole only and were susceptible to both voriconazole and posaconazole but with elevated MICs. Six isolates were resistant to voriconazole and four of them were also resistant to itraconazole, with the remaining two having intermediate resistance against itraconazole. Isolate L2731 was cross-resistant against itraconazole and posaconazole, whereas isolate L3131 was resistant to itraconazole and had intermediate resistance toward posaconazole. Interestingly, isolates from the same soil sample yielded different susceptibility results. Among the seven azole-susceptible isolates, six of them were confirmed to be susceptible to all three medical azoles. One isolate (F311), exhibited resistance to itraconazole and elevated MIC values for voriconazole and posaconazole, 0.25 g/ml and 0.125 g/ml, respectively.

**Table 2.** Characteristics of azole-resistant *A. fumigatus* isolates and the control isolates.

| Soil sample | Sampling site | Isolate | MIC ($\mu\text{g/ml}$)<sup>a</sup> | | | *cyp51A* mutation | *csp1* type |
| --- | --- | --- | --- | --- | --- | --- | --- |
|  |  |  | ITC | VRC | POS |  |  |
| L10 | Waterlow Park, London | L1031 | 2 | 8 | 0.0625 | TR<sub>46</sub>/Y121F/T239A | t02 |
|  |  | L1032 | 4 | 4 | 0.125 | TR<sub>46</sub>/Y121F/T239A | t02 |
|  |  | L1041 | 4 | 8 | 0.125 | TR<sub>46</sub>/Y121F/T239A | t02 |
|  |  | L1043 | 4 | 0.25 | 0.125 | TR<sub>34</sub>/L98H | t02 |
| L27 | Waterlow Park, London | L2731 | 4 | 1 | 0.5 | TR<sub>34</sub>/L98H | t02 |
|  |  | L2741 | 4 | 8 | 0.125 | TR<sub>46</sub>/Y121F/T239A | t09 |
| L28 | Whittington Hospital, London | L2831 | 4 | 8 | 0.125 | TR<sub>46</sub>/Y121F/T239A | t01 |
|  |  | L2841 | 4 | 0.25 | 0.125 | unknown<sup>b</sup> | t02 |
| L12 | Royal Free Hospital, London | L1231 | 4 | 0.5 | 0.125 | TR<sub>34</sub>/L98H | t02 |
|  |  | L1241 | 4 | 0.25 | 0.125 | TR<sub>34</sub>/L98H | t02 |
| L29 | Royal Free Hospital, London | L2931 | 4 | 0.125 | 0.0625 | TR<sub>34</sub>/L98H | t01 |
| L31 | Royal Free Hospital, London | L3131 | 4 | 1 | 0.25 | TR<sub>34</sub>/L98H | t02 |
|  |  | L3141 | 4 | 0.25 | 0.125 | TR<sub>34</sub>/L98H | t02 |
| L19 | Hyde Park, London | L1931 | 2 | 4 | 0.0625 | TR<sub>46</sub>/Y121F/T239A | t18 |
| L41 | Brompton Cemetery, London | L4131 | 4 | 0.25 | 0.125 | TR<sub>34</sub>/L98H | t02 |
| BS1 | Bath city center | BS131 | 4 | 0.25 | 0.125 | TR<sub>34</sub>/L98H | t02 |
| BS9 | Stonehenge, Wiltshire | BS941 | 4 | 0.125 | 0.0625 | TR<sub>34</sub>/L98H | t02 |
|  |  | BS942 | 4 | 0.125 | 0.125 | TR<sub>34</sub>/L98H | t02 |
| NF63 | New Forest, Hampshire | NF6341 | 4 | 0.25 | 0.125 | TR<sub>34</sub>/L98H | t04B |
| F3 | Elstree Open Space, London | F311 | 4 | 0.25 | 0.125 | TR<sub>34</sub>/L98H | t04B |
| F44 | Lavender farm, Surrey | F4411 | 0.125 | 0.031 | 0.016 | wild type | t05 |
| F55 | Wheat farm, Cambridge | F5511 | 0.25 | 0.063 | 0.031 | wild type | t04A |
| L11 | Royal Free Hospital, London | L1111 | 0.25 | 0.031 | 0.016 | wild type | t04A |
| L42 | Brompton Cemetery, London | L4211 | 0.125 | 0.031 | 0.031 | wild type | t03 |
| BS10 | Stonehenge, Wiltshire | BS1011 | 0.125 | 0.063 | 0.031 | wild type | t04A |
| NF42 | New Forest, Hampshire | NF4211 | 0.125 | 0.063 | 0.016 | wild type | t08 |
| wild-type control |  | Af293 | 0.25 | 0.063 | 0.016 | wild-type | t06 |
| azole-resistant control |  | BUU09 | 4 | 0.25 | 0.125 | TR<sub>34</sub>/L98H | t04B |

a. ITC, itraconazole; VRC, voriconazole; POS, posaconazole.
b. Isolate L2841 amplified poorly using the primers in this study.

### Molecular determination of the azole resistance mechanisms

The molecular mechanisms of azole resistance were determined by sequencing part of the *cyp51A* gene and its promoter region. Of the 19 azole-resistant isolates, 13 were found to be harbouring the TR_34_/L98H allele and six the TR_46_/Y121F/T289A allele (Table 2). All six azole susceptible isolates had the wild-type *cyp51A* allele. Both the negative and positive control isolates (Af293 and BUU09) carried the wild-type and TR_34_/L98H allele respectively.

### Molecular characterisation of *A. fumigatus*

Sequence typing of *csp1* yielded 9 different *csp1* types (Table 2). The six azole-susceptible wild-type environmental isolates were distributed over *csp1* subtypes t03 (1/6), t04A (3/6), t05 (1/6) and t08 (1/6). The 20 azole-resistant isolates were spread across five *csp1* subtypes, t01 (2/20), t02 (14/20), t04B (2/20), t09 (1/20) and t18 (1/20). All the TR_34_/L98H and TR_46_/Y121F/T289A isolates predominantly belonged to *csp1* subtype t02. The 13 isolates with the TR_34_/L98H allele were distributed over three *csp1* subtypes t01 (1/13), t02 (10/13) and t04B (2/13). The isolates carrying TR_46_/Y121F/T289A allele were grouped into four *csp1* subtypes: t01 (1/6), t02 (3/6), t09 (1/6) and t18 (1/6). The control isolate Af293, which was isolated from a patient with invasive aspergillosis belonged to *csp1* subtype t06 and the azole resistant control isolate BUU09 belonged to the t04B *csp1* subtype, which agreed with previously published results (24).

## Discussion

Given that AR*Af* continues to emerge globally, it is imperative that environmental monitoring, particularly in previously unsampled locations, is sustained. Here we present environmental sampling of London and the south east of England, home to approximately 15 million people living both urban and rural lifestyles. We show that azole-resistant *A. fumigatus* was detected in 6.7% (12/178) of the soil samples collected, and in 9.2% (12/131) of the soil samples positive for *A. fumigatus* growth. The prevalence of AR*Af* in the environment was 13.8% in urban areas (10/72), which included Greater London and Bath. In contrast, zero resistant isolates were found in soil samples collected from agricultural land and just two resistant isolates were found in rural samples collected from non-agricultural land (1.1%). We also report the UK’s first environmental AR*Af* isolates confirmed to be carrying the TR_46_/Y121F/T239A resistance allele, alongside an expanded distribution of the TR_34_/L98H allele.

Prevalence of AR*Af* in the UK was found to be lower than that of other European countries and Colombia (15, 17, 25–29), but higher than most Asian countries (24, 30–33), with the exception of India (29, 34). Our findings are also in close agreement with a recent UK-based environmental prevalence study in Wales, where AR*Af* was detected in 4.5% (30/671) of soil samples, with resistance predominantly found in urban city locations (13). However, they also describe a notable contrast to the only other UK based study, which found zero AR*Af* in urban locales but four resistant isolates from agricultural sites (1.7%) (14).

Our findings, and that of the Welsh study (13), appear to contradict the hypothesis that UK AR*Af* is driven by the environmental application of azoles in arable agriculture (35, 36). Indeed, of the 53 samples collected directly on or surrounding agricultural land, zero azole-tolerant isolates were identified. Rather, we found that the prevalence of resistance was higher in urban city centres, specifically flower beds and gardens, a finding that lends itself more readily to the hypothesis that the expanding range of AR*Af* stems more from the distribution and cultivation of horticultural crops, such as flowers, ornamentals and vegetables (15–17)

One particularly concerning discovery was the repeat isolation of AR*Af* genotypes – TR_34_/L98H and TR_46_/Y121F/T289A – from flower beds surrounding city centre hospitals. Owing to the opportunistic nature of *A. fumigatus*, its ability to cause debilitating illnesses in immunocompromised patients and the elevated mortality that is associated with IA caused by azole-resistant genotypes (22), the gardened areas around hospitals could be considered high risk locations if AR*Af* is present. Concern over the use of azole treated flower bulbs in hospital environments has been raised previously (15, 16, 37), and although we do not link any cases of azole-resistant aspergillosis to the isolates found in this study, our findings do add to a worrying trend of AR*Af* populated soil sampled from flower beds (15, 17).

The continued emergence of AR*Af* in urban city centres fits with the observation that most isolates belonged to a single *csp1* subtype (t02), a pattern consistent with the selective sweep of drug resistant genotypes (38). However, despite this obvious pattern of selection, city centres still harboured greater diversity than rural locations. All *csp1* subtypes identified during this study were found in Central London, suggesting an anthropological driver facilitating the migration of many *A. fumigatus* genotypes into a densely populated metropolis. Consequently, urban city centres, where *A. fumigatus* diversity is high, could facilitate a ‘melting pot’, where resistance alleles inadvertently brought into the city are given the opportunity to introgress onto novel genetic backgrounds via recombination, increasing the diversity potential of AR*Af*. Indeed, in London alone, TR_34_/L98H was found on two *csp1* subtypes and TR_46_/Y121F/T298A on four CSP subtypes.

Ultimately, this study highlights the importance of urban environments in the epidemiology of AR*Af*. We have shown that although AR*Af* appears to be environmental by origin, urban city centres, that are densely populated, are of high importance when mitigating strategies are considered. The use of azole treated bulbs in the environment around hospitals should be reconsidered and the continuous global monitoring of AR*Af* is imperative.

**Fig 1:**
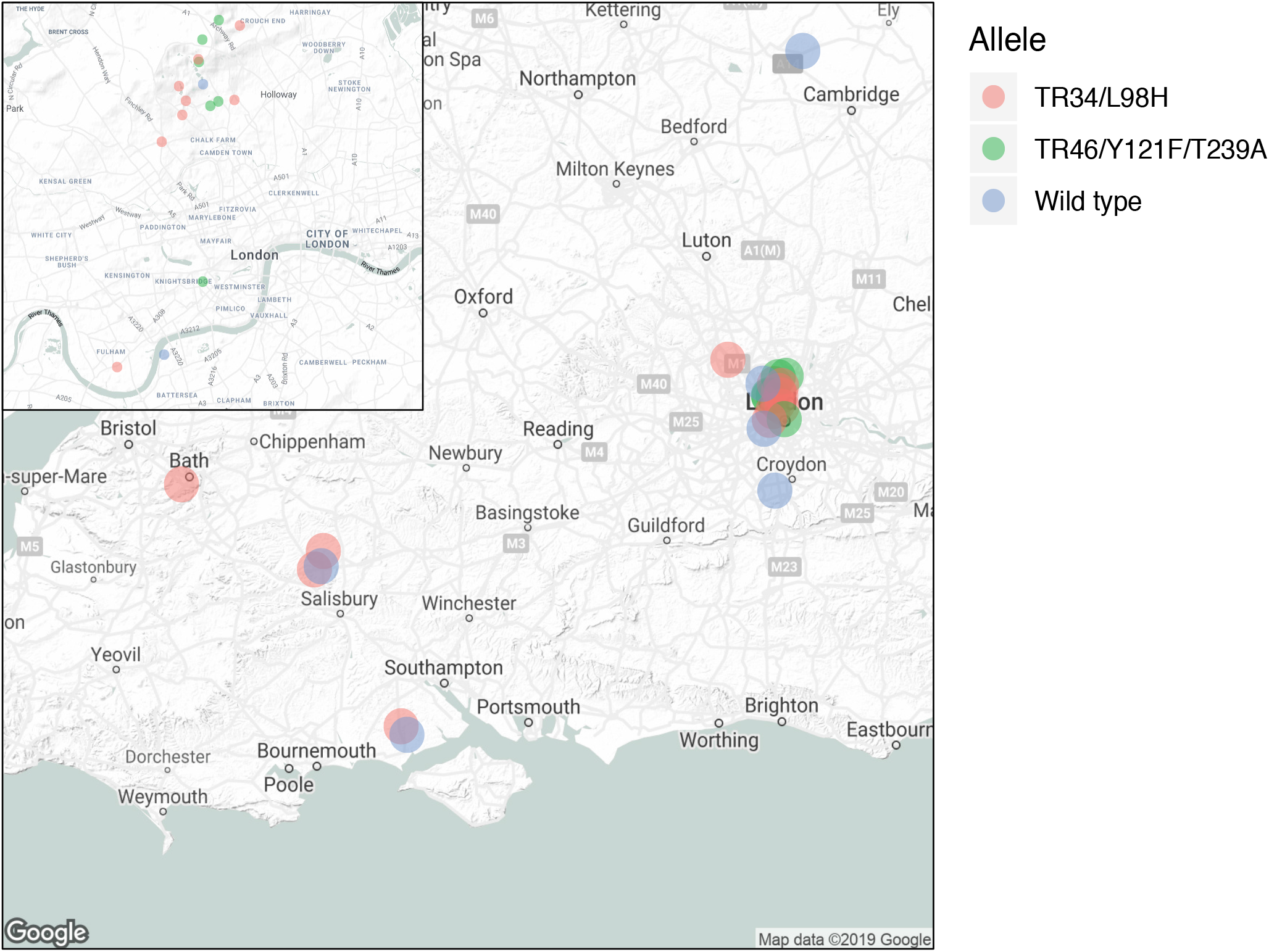
Distribution of azole susceptible and resistant *Aspergillus fumigatus* isolates collected in London (b) and southern UK counties (a). Isolates haboring cyp51a allele TR_34_/L98H are depicted in red, TR46/Y121F/T289A in blue and wild-type in green.

## Acknowledgments

The authors would like to thank Ali Abdolrasouli, Darius Armstrong-James and Andrew Scourfield for their helpful discussions whilst analysing the data. We also acknowledge joint Centre funding from the UK Medical Research Council and Department for International Development. Funding: T.R.S., A.P.B., J.R. and M.C.F. were supported by the Natural Environmental Research Council (NERC; NE/P001165/1) and all authors were supported by the Medical Research Council (MRC; MR/R015600/1).

## Author Contributions

T.R.S., Y.Z. and M.C.F. conceived and designed the study. T.R.S., Y.Z., A.P.B. and J.M.G.S. collected the data. T.R.S., Y.Z. and A.P.B. analyzed the data. T.R.S. and Y.J. wrote the manuscript. T.R.S., Y.Z., A.P.B., J.M.G.S., J.R., and M.C.F. discussed the results and commented on the manuscript

